# *metagenomeFeatures*: An R package for working with 16S rRNA reference databases and marker-gene survey feature data

**DOI:** 10.1101/339812

**Authors:** Nathan D. Olson, Nidhi Shah, Jayaram Kancherla, Justin Wagner, Joseph N. Paulson, Hector Corrada-Bravo

## Abstract

We developed the *metagenomeFeatures* R Bioconductor package along with annotation packages for the three primary 16S rRNA databases (Greengenes, RDP, and SILVA) to facilitate working with 16S rRNA sequence databases and marker-gene survey feature data. The *metagenomeFeatures* package defines two classes, `MgDb` for working with 16S rRNA sequence databases, and `mgFeatures` for working with marker-gene survey feature data. The associated annotation packages provide a consistent interface to the different 16S rRNA databases facilitating database comparison and exploration. The `mgFeatures` represents a crucial step in the development of a common data structure for working with 16S marker-gene survey data in R.

**Availability:** https://bioconductor.org/packages/release/bioc/html/metagenomeFeatures.html

## 1 Introduction

16S rRNA marker-gene surveys have significantly advanced our understanding of the diversity and structure of prokaryotic communities present in ecosystems including the human gut, open ocean, and even the international space station (Lang *et al*., 2017; Thompson *et al*., 2017; Human Microbiome Project Consortium 2012). For a 16S rRNA marker-gene survey, the 16S rRNA gene is sequenced using a targeted assay. The raw sequence data is processed using a bioinformatic pipeline where the sequences are grouped into features, e.g., operational taxonomic units (OTUs) or sequence variants (SVs), yielding a set of representative sequences (Callahan, McMurdie, and Holmes 2017; Beiko 2015).

A critical step in 16S rRNA marker-gene surveys is comparing representative sequences to a reference database for taxonomic classification or phylogenetic placement (Nguyen *et al*., 2016). There are numerous 16S rRNA reference databases of which Greengenes, RDP, and SILVA are arguably the most commonly used (DeSantis et al. 2006; Cole et al. 2014; Quast et al. 2012; McDonald et al. 2012). Additionally, there are smaller system-specific databases such as HOMD for the human oral microbiome (Chen et al. 2010)(http://www.homd.org/) and soil reference database (Choi et al. 2017). System-specific databases can improve taxonomic assignments for microbial communities not well represented in the major databases (Rohwer et al. 2017).

16S rRNA databases differ in the number and diversity of sequences, the taxonomic classification system, and the inclusion of intermediate ranks (Balvočiūtė and Huson 2017, Table 1). Databases format their data differently and use sequence identification systems unique to their database, challenging membership and composition comparisons. For example, Yang et al. (2016) used the SILVA database to evaluate how different 16S rRNA variable regions impact phylogenetic analysis. Similarly, Martinez-Porchas et al. (2017) also used the SILVA database to evaluate sequence similarity between 16S rRNA gene conserved regions. Differences in database formatting present a significant barrier to performing the same analysis using multiple databases. Additionally, taxonomic assignments can be database-dependent, providing further justification for database comparisons (Pettengill and Rand 2017). To facilitate database comparisons RNACentral (http://rnacentral.org/) a resource combining non-coding RNA databases, provides unique identifiers for the sequences (The RNAcentral Consortium 2017).

**Table 1.** 16S rRNA gene sequence databases with Bioconductor annotation packages we developed.

| Database | Version | Sequences | Taxonomic System<sup>1</sup> |
| --- | --- | --- | --- |
| Greengenes | 13.5<sup>2</sup> | 1,262,986 | NCBI |
| RDP | 11.5 | 3,356,809 | Bergey's |
| SILVA | 128.1 | 1,922,213 | Bergey's |

<sup>1</sup> Primary taxonomic system see (see Balvočiūtė and Huson 2017) and references therein for additional information on databases.
<sup>2</sup> Greengenes 13.8 85% OTUs is included in the *metagenomeFeatures* package as an example `MgDb` formatted database.

The statistical programming language, R provides a rich environment and software for data analysis (R Core Team, n.d.). Additionally, Bioconductor, the R bioinformatic software resource (Huber et al. 2015) includes a number of packages for working with DNA sequence data and 16S rRNA marker-gene survey data such as *phyloseq* (McMurdie and Holmes 2013) and *metagenomeSeq* (Paulson et al. 2013). While a number of software tools are available for working with 16S rRNA marker-gene survey feature data including Mothur (Schloss et al. 2009) as well as QIIME, specifically the q2-feature-classifıer plugin (Bokulich et al. 2018). There are no existing tools for working with multiple 16S rRNA databases. Furthermore, tools for working with 16S rRNA marker-gene survey feature data in R all use different data structures. Therefore, an R package defining consistent data structures for working with 16S rRNA database and marker-gene survey feature data is needed.

To address this need we developed the R package *metagenomeFeatures* for working with both 16S rRNA gene database and marker-gene survey feature data. *metagenomeFeatures* provides a common data structure for working with the 16S rRNA databases and marker-gene survey feature data. Additionally, this package is the first step towards the development of a common data structure for use in analyzing metagenomic and marker-gene survey data using R packages such as *phyloseq* (McMurdie and Holmes 2013) and *metagenomeSeq* (Paulson et al. 2013).

## 2 *metagenomeFeatures* Package

The *metagenomeFeatures* package defines two data structures, `MgDb` for working with 16S rRNA databases, and `mgFeatures` for working with marker-gene survey feature data. There are three types of relevant information for both `MgDb` and `mgFeatures` class objects, (1) the sequences themselves, (2) sequence taxonomic lineage, and (3) a phylogenetic tree representing the evolutionary relationship between features. `MgDb` and `mgFeatures` data structures are both S4 object-oriented classes with slots for taxonomy, sequences, phylogenetic tree, and metadata.

The `MgDb-class` provides a consistent data structure for working with different 16S rRNA databases. As shown in Table 1, 16S rRNA databases contain hundreds of thousands to millions of sequences, therefore an SQLite database is used to store the taxonomic and sequence data. Using an SQLite database prevents the user from loading the full database into memory. The database connection is managed using the *RSQLite* R package (Müller et al. 2017), and the taxonomic data are accessed using the *dplyr* and *dbplyr* packages (Wickham 2017; Wickham et al. 2017). The *DECIPHER* package is used to format the sequence data as an SQLite database (Wright 2016) and provides functions for working directly with the sequence data in the SQLite database. The `phylo` class, from the *APE* R package, defines the tree slot (Paradis, Claude, and Strimmer 2004). We developed Bioconductor annotation packages for commonly used databases, Greengenes, RDP, and SILVA (Table 1, Cole et al. 2014; Quast et al. 2012; DeSantis et al. 2006). Along with database specific sequence identifiers, RNAcentral indentifers are included in the SQLite table for inter-database comparisons.

`mgFeatures-class` is used for storing and working with marker-gene survey feature data. Similar to the `MgDb-class`, the `mgFeatures-class` has four slots, for taxonomy, sequences, phylogenetic tree, and metadata. As the number of features in a marker-gene survey dataset is significantly smaller than the number of sequences in a reference database, `mgFeatures` uses common Bioconductor data structures, `DataFrames` and `DNAStringSets` to define the taxonomic and sequence slots (Pagès et al. 2008; Pagès, Lawrence, and Aboyoun 2017). Similar to `MgDb-class`, a `phylo` class object is used to define the tree slot. For both `MgDb` and `mgFeatures` classes the tree slot is optional and the metadata are stored as a list.

The *metagenomeFeatures* package includes vignettes as example use cases for the *metagenomeFeatures* package and associated reference database annotation packages.

- Retrieving sequence and phylogenetic data for OTUs from closed-reference clustering.
- Exploring diversity for a taxonomic group of interest.

The R command `browseVignettes` (“`metagenomeFeatures`”) provides a list of vignettes associated with the package and vignette(“x”) is used to view specific `vignettes`, where “`x`” is the vignette name.

To further demonstrate the utility of the package, the manuscript supplemental information demonstrates using *metagenomeFeatures, greengenes13.5MgDb* annotation package, and *DECIPHER* to evaluate the potential for species-level taxonomic classification using 16S rRNA V12 and V4 sequence data.

## 3 Conclusions

The *metagenomeFeatures* package provides data structures and functions for working with 16S rRNA gene sequence reference databases and marker-gene survey feature data. The data structure provided by the `MgDb-class` in conjunction with the shared sequence identifier system developed by RNACentral facilitates comparisons between 16S rRNA databases. The `mgFeatures-class` provides the groundwork for the development of a common data structure for working with metagenomic and marker-gene sequence data in R which will increase interoperability between R packages developed for working with metagenomic sequence data. Additionally, while the data structures were developed for 16S rRNA gene sequence data they can be used for any marker-gene sequence data without modification and can be extended to work with shotgun metagenomic sequence data and databases.

## 4 Acknowledgements

The authors would like to thank Drs. Mihai Pop, Marc Salit, Samuel Forry, and Arlin Stolzfus for feedback on the manuscript. The Bioconductor core team provided valuable feedback during the package submission and update process. Opinions expressed in this paper are the authors’ and do not necessarily reflect the policies and views of NIST or affiliated venues. Certain commercial equipment, instruments, or materials are identified in this paper in order to specify the experimental procedure adequately. Such identification is not intended to imply recommendations or endorsement by NIST, nor is it intended to imply that the materials or equipment identified are necessarily the best available for the purpose. Official contribution of NIST; not subject to copyrights in USA.

